# Sparse and Skew Hashing of K-Mers

**DOI:** 10.1101/2022.01.15.476199

**Authors:** Giulio Ermanno Pibiri

## Abstract

**Motivation:** A dictionary of *k*-mers is a data structure that stores a set of *n* distinct *k*-mers and supports membership queries. This data structure is at the hearth of many important tasks in computational biology. High-throughput sequencing of DNA can produce very large *k*-mer sets, in the size of billions of strings – in such cases, the memory consumption and query efficiency of the data structure is a concrete challenge.

**Results:** To tackle this problem, we describe a *compressed* and *associative* dictionary for *k*-mers, that is: a data structure where strings are represented in compact form and each of them is associated to a unique integer identifier in the range [0, *n*). We show that some statistical properties of *k*-mer *minimizers* can be exploited by minimal perfect hashing to substantially improve the space/time trade-off of the dictionary compared to the best-known solutions.

**Availability:** The C++ implementation of the dictionary is available at https://github.com/jermp/sshash.

## 1 Introduction

A *k*-mer is a string of length *k* over the DNA alphabet {A, C, G, T}. Software tools based on *k*-mers are in widespread use in Bioinformatics. Many large-scale analyses of DNA share the elementary need of determining the exact membership of *k*-mers to a given set S, i.e., they rely on the space/time efficiency of a *dictionary* data structure for *k*-mers [Chikhi et al., 2021]. This work proposes such an efficient dictionary. More precisely, the problem we study here is defined as follows. Given a large string over the DNA alphabet (e.g., a genome or a pan-genome), let 𝒮 be the set of *all* its distinct *k*-mers, with | 𝒮 | = *n*. A dictionary for 𝒮 is a data structure that supports the following two operations:

- for any *k*-mer *g*, Lookup(*g*) returns a unique integer 0 ≤ *i* < *n* if *g* ∈ 𝒮 (*i* is the “identifier” of *g* in 𝒮) or *i* = −1 if *g* < 𝒮;
- for any 0 ≤ *i* < *n*, Access(*i*) extracts the *k*-mer *g* for which Lookup(*g*) = *i*.

By means of the Lookup query, the dictionary is able to answer membership queries in an exact way (rather than approximate) and to associate satellite information to *k*-mers (such reference identifiers or abundances). Thanks to the Access query, the original set 𝒮 can be reconstructed, meaning that the dictionary is a self-index for 𝒮.

In sequence analysis tasks, it is very often the case that we are given a pattern *P* of length | *P* | ≥ *k* and we are interested in answering membership to 𝒮 for all the *k*-mers read *consecutively* from *P*, that is, for *P*[*i, i* + *k*), *i* = 0, …, | *P* |−*k*. For example, we may decide that the whole pattern *P* is present in a genome if the number of *k*-mers of *P* that belong to 𝒮 is at least *θ* · (| *P* | − *k* + 1), for a prescribed coverage threshold *θ* > 0, such as *θ* = 0.8 [Solomon and Kingsford, 2016, Bingmann et al., 2019]. In other words, Lookup queries are often issued for *consecutive k*-mers (one being the previous shifted to the right by one symbol) [Robidou and Peterlongo, 2021]. While it is obviously possible to perform | *P* | − *k* + 1 Lookup queries for a pattern of length | *P* |, it also seems profitable to answer “Is *P*[*i, i* + *k*) a member of S?” more efficiently knowing that the previous *k*-mer shares *k* − 1 symbols with *P*[*i, i* + *k*). We regard this latter scenario as that of *streaming* queries.

Therefore, our objective is to support Lookup, Access, and streaming membership queries as efficiently as possible in compressed space. (The data structure is static: insertions/deletions of *k*-mers are not supported.)

As a first introductory remark we shall mention that the algorithmic literature about the so-called *compressed string dictionary* problem is rich of solutions, e.g., based on Front-Coding, tries, hashing, or combinations of such techniques (see the survey by Martínez-Prieto et al. [2016]). However, these solutions are unlikely to be competitive for the specialized version of the problem we tackle here because they are relevant for “generic” strings that usually: (1) have variable length; (2) are drawn from larger alphabets (e.g., ASCII); (3) do not exhibit particular properties that can aid compression. Instead, *k*-mers are fixed-length strings; their alphabet of representation is very small (just 2 bits per alphabet symbol are sufficient); and since *k*-mers are extracted consecutively from DNA, two consecutive strings overlap by *k* − 1 symbols that are redundant and should *not* be represented twice in the dictionary. This motivates the study of specialized solutions for *k*-mers.

These properties are elegantly captured by the *de Bruijn graph* representation of 𝒮 – a graph whose nodes are the *k*-mers in 𝒮 and the edges model the string overlaps between the *k*-mers. Using this formalism, it is possible to reduce the redundancy of the symbols in S by considering *paths* in the graph and their corresponding strings. We will better formalize this point in Section 2.

For the scope of this work, it is sufficient to point out that: (1) many algorithms have been proposed to build de Bruijn graphs efficiently [Chikhi et al., 2016, Khan and Patro, 2021, Khan et al., 2021] from which these paths can be extracted for indexing purposes; (2) not surprisingly, essentially all state-of-the-art dictionaries for *k*-mers – that we briefly review in Section 3 – are based on the principle of indexing such collections of paths [Chikhi et al., 2014, Almodaresi et al., 2018, Rahman and Medvedev, 2020, Marchet et al., 2021]. We also follow this direction.

However, we note that existing dictionary data structures either represent such paths with an FM-index [Ferragina and Manzini, 2000] (or one of its many variants), hence retain highly compressed space but very slow query time in practice or, vice versa, resort on hashing for fast evaluation but take much more space [Almodaresi et al., 2018, Marchet et al., 2021]. It is, therefore, desirable to have a good balance between these two extremes.

For this reason, we show that we can still enjoy the query efficiency of hashing while taking small space – significantly less space compared to existing schemes also based on hashing. More specifically, we show how two statistical properties of *k*-mer *minimizers* – those of being *sparse* and *skewly distributed* in DNA sequences – can be better exploited to derive an efficient dictionary based on minimal perfect hashing and compact encodings (Section 4). We evaluate the proposed data structure over sets of billions of *k*-mers, under different query distributions and modalities, and exhibit a substantial performance improvement compared to prior solutions for the problem (Section 5). Our C++ implementation of the dictionary is available at https://github.com/jermp/sshash.

## 2 Preliminaries

In this section we give some preliminary remarks to better support the exposition in Section 3 and 4. Let 𝒮 be the collection of the *n* distinct *k*-mers extracted from a given, large, string (or set of strings). This string can be, for example, the genome of an organism.

Throughout the paper, we consider to be identical two *k*-mers that are the *reverse complement* of each other.

### Path Covering the de Bruijn Graph

A de Bruijn graph (dBG) for S is a directed graph *G*(S) where the set of nodes is S and a direct edge from node *u* to *v* exists if and only if the last *k* − 1 symbols of *u* are equal to the first *k* − 1 symbols of *v*, i.e., *u*[1, *k* − 1] = *v*[0, *k* − 2]. It follows that a path traversing 𝓁 nodes corresponds to a string of length 𝓁 + *k* − 1 spelled out by the path, obtained by “glueing” all the nodes’ *k*-mers in order.

A disjoint-node path cover for *G*(S) is a set of paths where *each* node belongs to *exactly one* path, e.g., a set of unitigs, maximal unitigs, or maximal *stitched* unitigs [Rahman and Medvedev, 2020] (also known as *simplitigs* [Břrinda et al., 2021]). We denote such a cover with 𝒮 ′.

The strings in 𝒮′ form the natural basis for a space-efficient dictionary because: (1) by considering paths in the graph, the number of symbols in 𝒮′ is less than the number of symbols in the original 𝒮 and, (2) by being a disjoint-node path cover we are guaranteed that there are *no duplicate k*-mers in 𝒮′. Therefore, we assume from now on that a path cover 𝒮′ has been computed for *G*(𝒮) as the input for our problem.

### Minimizers and Super-*k*-mers

Given a *k*-mer *g*, an integer *m* ≤ *k*, and a total order relation *R* on all *m*-length strings, the smallest *m*-mer of *g* according to *R* is called the *minimizer* of *g. R* could be, for example, the simple lexicographic order. Instead, here we use the random order given by a hash function *h*, chosen from a universal family. Therefore, simply put, the minimizer of *g* is the *m*-mer of *g* that minimizes the value of *h* (sometimes called a “random” minimizer).

Minimizers are very popular in sequence analysis, such as for seed-and-extend algorithms, because of the following empirical property: consecutive *k*-mers tend to have the same minimizer [Schleimer et al., 2003, Roberts et al., 2004]. This means that there are far less distinct minimizers than *k*-mers – approximately, (*k* − *m* + 2)/2 times less minimizers than *k*-mers (independently of the sequence length), if *m* is not very small compared to *k* (more precisely, see Zheng et al. [2020, Theorem 3]). For example, if *k* = 31 and *m* = 20, we should expect to see ≈ 6.5× less minimizers than *k*-mers.

Given a string *S* of length at least *k* (e.g., a path in a dBG), we call a *super-k-mer* of 𝒮 a maximal sequence of consecutive *k*-mers having the same minimizer [Li et al., 2013].

### Minimal Perfect Hashing

Given a set 𝒳 of *n* distinct keys, a function *f* that maps bijectively the keys into the integer range {0, …, *n* − 1} is called a minimal perfect hash function (MPHF) for the set 𝒳. The function is allowed to return an arbitrary value in [0, *n*) for any key that does not belong to 𝒳, hence it can be realized in small space, in practice 2 − 3 bits/key (albeit log_2_ *𝓁* ≈ 1.44 bits/key are sufficient in theory [Mehlhorn, 1982]). Many efficient algorithms have been proposed to build MPHFs from static sets that scale well to large values of *n* and retain practically-constant evaluation time. In this paper, we use PTHash [Pibiri and Trani, 2021a,b] for its very fast evaluation time, usually 2 − 4× better than other techniques, and good space effectiveness.

### Elias-Fano Encoding

Given a monotone integer sequence *S*[0..*n*) whose largest (last) element is less than or equal to a known quantity *U*, the Elias-Fano encoding represents *S* in at most *n* ⌈ log_2_(*U*/*n*) ⌉ + 2*n* bits [Fano, 1971, Elias, 1974]. With *o*(*n*) extra bits it is possible to decode any *S*[*i*] in constant time and support successor queries in *O*(log(*U*/*n*)) time. We point the interested reader to the survey by Pibiri and Venturini [2021, Section 3.4] for a complete description and discussion of the encoding.

Elias-Fano has been recently used as a key ingredient of many compressed, practical, data structures (see, e.g., [Pibiri and Venturini, 2017, 2019, Perego et al., 2021]).

## 3 Related Work

As anticipated in Section 1, most existing solutions for exact membership queries are based on indexing paths of the de Bruijn graph (see also Section 2), such as its unitigs or maximal (possibly, stitched) unitigs. These approaches have also been summarized in the recent survey by Chikhi et al. [2021, Section 4.2], hence we give a rather cursory overview here.

The paths can be represented using an FM-index [Ferragina and Manzini, 2000] for very compact space [Chikhi et al., 2014, Rahman and Medvedev, 2020]. The practical efficiency of the FM-index mainly depends on how many samples of the suffix-array are kept in the index.

Other approaches resort on hashing for fast lookup queries. For example, Bifrost [Holley and Melsted, 2020] uses a hash table of minimizers whose values are the locations of the minimizers in the unitigs. The index was designed to be dynamic, hence allowing insertion/removal of *k*-mers and consequent re-computation of the unitigs. The dynamic nature of Bifrost makes it consume higher space compared to static approaches using compressed hash representations and succinct data structures, like Pufferfish [Almodaresi et al., 2018] and Blight [Marchet et al., 2021]. Hence, it is regarded as out of scope for this work.

Pufferfish [Almodaresi et al., 2018] associates to each *k*-mer its location in the unitigs using a MPHF and a vector of absolute positions. The authors also propose a sparse version of the index where the vector of positions is sampled to improve space usage at the expense of query time. Blight [Marchet et al., 2021] is another associative dictionary based on minimal perfect hashing. All the super-*k*-mers having the same minimizer are grouped together into an index partition and a separate MPHF is built for all the *k*-mers in the partition. Since the *k*-mers’ offsets are relative to a given partition, the space usage is improved compared to Pufferfish. To further reduce space, a *k*-mer is associated to the segment of 2^*b*^ super-*k*-mers where it belongs to, for a given *b* ≥ 0. This reduces the space of the dictionary by *b* bits per *k*-mer but a lookup needs to scan (at most) 2^*b*^ super-*k*-mers. Very importantly, Pufferfish and Blight are also optimized for streaming membership queries.

## 4 Sparse and Skew Hashing

In this section we describe our main contribution: an exact, associative, and compressed dictionary data structure for *k*-mers, supporting fast Lookup, Access, and streaming queries. From a high-level point of view, the dictionary is obtained via a careful combination of minimal perfect hashing and compact encodings. In particular, we show how two important properties of minimizers – those of being sparse (Section 4.1) and skewly distributed (Section 4.2) in DNA strings – can be exploited to achieve an efficient dictionary. We aim at a good trade-off between dictionary space and query efficiency.

Recall from Section 2 that the dictionary is built from a collection of paths covering a de Bruijn graph (for example, the maximal stitched unitigs), that is: a collection of strings, each of length at least *k* symbols, with no duplicate *k*-mers. For ease of notation, we indicate with *p* the number of paths in the collection and with *N* their cumulative length (the total number of DNA bases in the input). The number of (distinct) *k*-mers is, therefore, *n* = *N* − *p*(*k* − 1).

### 4.1 Sparse Hashing

The starting point for our development is based on the well-known empirical property of minimizers in that consecutive *k*-mers are likely to have the same minimizer. Thus, instead of working with individual *k*-mers, we focus on maximal sequences of *k*-mers having the same minimizer – the so-called super-*k*-mers (see Section 2). Super-*k*-mers are useful because of the following two reasons.

- As super-*k*-mers are likely to span several consecutive *k*-mers, we expect to see far fewer super-*k*-mers than *k*-mers – approximately, (*k* − *m*+2)/2 times less for a large-enough minimizer length *m*. Informally, this property allows a space usage proportional to the number of super-*k*-mers, thus *sparsifying* the dictionary.
- A super-*k*-mer of length *K* is a space-efficient representation for its constituent *K* − *k* + 1 *k*-mers since it takes 2*K* /(*K* − *k* + 1) bits/*k*-mer instead of the trivial cost of 2*k* bits/*k*-mer.

Therefore, our refined ambition is to index the super-*k*-mers of the input using minimizers. Although this can simply be achieved via hashing the minimizers, that is, by concatenating all the super-*k*-mers having the same minimizer (like in the Blight index [Marchet et al., 2021]) – we claim that his approach is very wasteful in terms of space. In fact, note that each super-*k*-mer has a fixed cost of 2(*k* − 1) bits for representing the “tail” of its string (its last *k* − 1 symbols). This fixed cost is only well amortized (say, negligibly small) when the length of the super-*k*-mer is much larger than *k* − 1. In other words, when the super-*k*-mer contains many more *k*-mers than *k* − 1. While possible in some extreme cases (e.g., the same minimizer repeats in sequence), it is not usually so for the values of *k* and *m* used in concrete applications; actually, a super-*k*-mer is more likely to contain *k* − *m* + 1 *k*-mers or less.

If *z* indicates the number of super-*k*-mers in the input, then the space of this simple solution would be, at least, 2 + 2*z*(*k* − 1)/*n* bits/*k*-mer (extra space is then needed to accelerate the queries). For example, consider the whole human genome with *k* = 31 and *m* = 20. There are more than *z* = 396 × 10^6^ super-*k*-mers for, roughly, *n* = 2.5 × 10^9^ distinct *k*-mers. Therefore, partitioning the strings according to super-*k*-mers would cost at least 11.50 bits/*k*-mer. As we will better see in Section 5, our dictionary can be tuned to take, *overall*, 8.28 bits/*k*-mer in this case (or less).

Thus, it is of utmost importance to *not* break the strings according to super-*k*-mers if space-efficiency is a concern. Instead, we identify a super-*k*-mer in the strings, whose total length is *N*, with an absolute offset of ⌈ log_2_(*N*) ⌉ bits. To be precise, an offset is the position in [0, *N*) of the first base of a super-*k*-mer. Since *k* should be chosen large enough to allow good *k*-mer specificity, 2(*k* − 1) will be much larger than ⌈log_2_(*N*) ⌉ in practice, even for the largest genomes. For example, we use *k* = 31 in our experiments, as done in many other works [Almodaresi et al., 2018, Marchet et al., 2021, Rahman and Medvedev, 2020, Bingmann et al., 2019], whereas ⌈ log_2_(*N*) ⌉ is around 30 − 33 for collections with billions of *k*-mers (see also Table 2 at page 6). The use of absolute offsets can almost halve the space overhead for the indexing of super-*k*-mers in such cases. The space saving is even larger for larger *k*.

#### Dictionary Layout and Compression

Based on the above discussion, we now detail the different components of our dictionary data structure.

- *Strings*. The *p* strings in the input are written one after the other in a vector of 2*N* bits (2 bits per input base). We also keep in a sorted integer sequence of length *p* the endpoints of the strings to avoid detection of alien *k*-mers. This sequence, *Endpoints*, is compressed with Elias-Fano and takes *p* ⌈ log_2_(*N* /*p*) ⌉ + 2*p* + *o*(*p*) bits.
- *Minimizers*. Let ℳ be the set of all distinct minimizers seen in the input, with *M* = | ℳ |, and *z* the number of super-*k*-mers. Clearly, we have *z* ≥ *M* because a minimizer can appear more than once in the input. Given a minimizer *r*, let us call the *bucket* of the minimizer *r, B*_*r*_, the set of all the super-*k*-mers that have minimizer *r*. We build a minimal perfect hash function *f* for ℳ. The MPHF provides us an addressable space of size *M*: for a minimizer *r*, the value *f* (*r*) ∈ [0, *M*) is the “bucket identifier” of *r*. We keep an array *Sizes*[0, *M* +1), where *Sizes*[ *f* (*r*) + 1] = |*B*_*r*_ | is the size of the bucket of *r*, and *Sizes*[0] = 0. We then take the prefix-sums of *Sizes*, i.e., we replace *Sizes*[*i*] with *Sizes*[*i*] + *Sizes*[*i* − 1] for all *i* > 0. Therefore, for a given minimizer *r*, now *Sizes*[ *f* (*r*)] indicates that there are *Sizes*[ *f* (*r*)] super-*k*-mers before bucket *B*_*r*_ in the order given by *f*.

The MPHF costs roughly 3 bits per minimizer; the *Sizes* array is compressed with Elias-Fano too and takes (*M* +1) ⌈log_2_(*z*/(*M* +1)) ⌉ + 2(*M* + 1) + *o*(*M* + 1) bits.

- *Offsets*. The absolute offsets of the super-*k*-mers into the strings are stored in an array, *Offsets*[0, *z*), in the order given by *f*. For a minimizer *r* such that *Sizes*[ *f* (*r*)] = *begin*, its |*B*_*r*_ | offsets are written consecutively (and in sorted order) in *Offsets*[*begin, begin* + |*B*_*r*_ |). Note that, by construction, *Sizes*[ *f* (*r*) + 1] − *Sizes*[ *f* (*r*)] = |*B*_*r*_ | > 0. The space for the *Offsets* array is *z* ⌈ log_2_(*N*) ⌉ bits.

Fig. 1 illustrates the different components of the dictionary and provides a concrete example for an input collection of 4 strings. Next we describe how the Lookup and Access queries are supported.

**Fig. 1.**
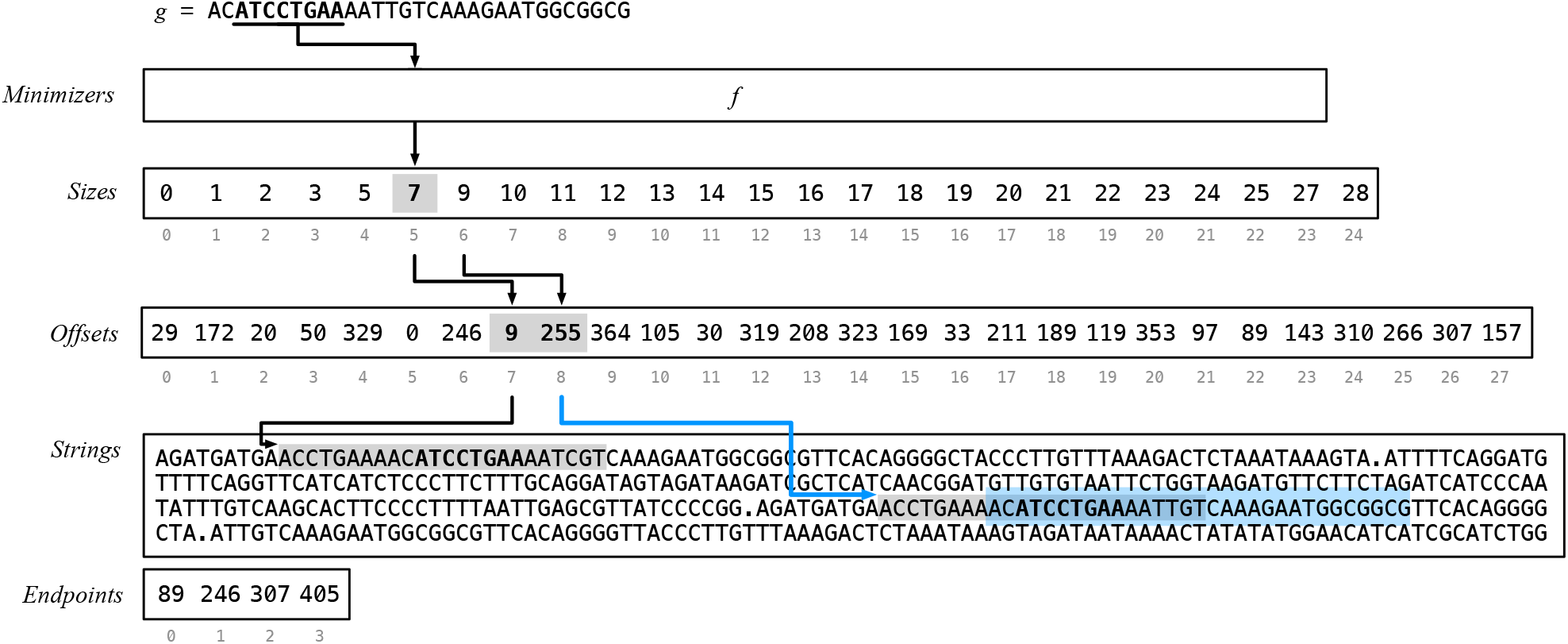
A schematic representation of the proposed dictionary data structure. The input of the example contains *p* = 4 strings (pictorially separated by a “.” symbol, but practically by the *Endpoints* array) for a total of *N* = 405 bases, and *N* − *p*(*k* − 1) = 405 − 4 · (31 − 1) = 285 *k*-mers for *k* = 31. There are *M* = 24 minimizers for *m* = 8 and *z* = 28 super-*k*-mers, thus the *Sizes* and *Offsets* arrays have length, respectively, *M* + 1 = 25 and *z* = 28. All minimizers have bucket size equal to 1 except for 3 of them (i.e., AACCTGAA, ATCCTGAA, TGTCAAAG) that have bucket size equal to 2. The picture also shows an example of Lookup for the *k*-mer *g* = AC**ATCCTGAA**AATTGTCAAAGAATGGCGGCG, whose minimizer *r* = ATCCTGAA is highlighted in bold font. The flow of the algorithm is represented by the arrows. First, the function *f* returns the identifier of *r* as *f* (*r*) = 5. Then the bucket size of *r* is computed: in this case, we have |*Br* | = *end* − *begin* = *Sizes*[5 + 1] − *Sizes*[5] = 9 − 7 = 2, indicating that there are 2 super-*k*-mers to consider. The offsets of the super-*k*-mers are retrieved as *Offsets*[*begin*] = *Offsets*[7] = 9 and *Offsets*[*begin* + 1] = *Offsets*[8] = 255. The two super-*k*-mers are scanned in *Strings* starting at *Strings*[9] and *Strings*[255] respectively. (At most *k* − *m* + 1 = 31 − 8 + 1 = 24 *k*-mers are considered in each super-*k*-mer, as highlighted by the gray box. See the Supplementary Material for a discussion about this point.) Lastly, the *k*-mer *g* is found at position *w* = 8 in the second super-*k*-mer, that is, at offset *t* + *w* = 255 + 8 = 263. Since there are two strings before the one containing *g*, then *j* = 2, and we have to discard *j*(*k* − 1) = 2 · (31 − 1) = 60 invalid ranks for the calculation of the identifier *i* of *g*. Therefore, we return *i* = *t* + *w* − *j*(*k* − 1) = 263 − 60 = 203.

#### Lookup

We first recall that the Lookup query takes as input a *k*-mer *g* and returns a unique identifier *i* for *g*: *i* ∈ [0, *n*) if *g* is found in the dictionary, or *i* = −1 otherwise. The Lookup algorithm is as follows.

We compute the minimizer *r* of *g* and its bucket identifier as *f* (*r*). Then, we locate the super-*k*-mers in its bucket *B*_*r*_ by retrieving the corresponding offsets from *Offsets*[*begin, end*), where *begin* = *Sizes*[ *f* (*r*)] and *end* = *Sizes*[ *f* (*r*) + 1]. For every offset *t* in *Offsets*[*begin, end*), we scan the super-*k*-mer starting from *Strings*[*t*] comparing its *k*-mers to the query *g*. If *g* is not found, we just return −1. Instead, if *g* is found in position *w* in the super-*k*-mer, we return the “identifier” *i* of *g* as *i* = *t* + *w* − *j*(*k* − 1), where *j* < *p* is the number of strings *before* the one containing the offset *t* (this quantity is computed from the *Endpoints* array).

Refer to Fig. 1 for an example of Lookup and to the Supplementary Material for further technical details.

#### Double Strandedness

A detail of crucial importance for the Lookup algorithm is *double strandedness*. A *k*-mer and its reverse complement are considered to be identical. This means that if a *k*-mer *g* is not found by the Lookup algorithm, there can still be the possibility for its reverse complement *ĝ* to be found in *Strings*. Therefore, the actual Lookup routine will first search for *g* and – only if not found – will also search for *ĝ*. This effectively doubles the query time for Lookup in the worst case.

To guarantee that a Lookup will always inspect *one* single bucket, we use a different minimizer computation (during both query and dictionary construction): we select as minimizer the minimum between the minimizer of *g* and that of *ĝ*. In this way, it is guaranteed that two *k*-mers being the reverse complements of each other always belong to the same bucket.

This different minimizer selection actually changes the parsing of super-*k*-mers from the input during the construction of the dictionary. We refer to this parsing modality as *canonical* henceforth, in contrast to the *regular* modality we assumed so far. When this modality is chosen, we expect to see an increase in the number of distinct minimizers used (on average, the minimizers of *g* and *ĝ* have equal probability of being the minimum one) for a higher space usage, but faster query time.

We will explore the space/time trade-off between the regular and canonical modalities in Section 5.1.

#### Access

The Access query retrieves the *k*-mer string *g* given its identifier *i*. This identifier represents the rank of *g* in *Strings* but, since the strings have variable lengths and their last *k* − 1 symbols do not correspond to valid ranks, we cannot directly access *Strings* at position *i*. Instead, we have to compute the offset *t* corresponding to the *k*-mer *g* of rank *i*. Therefore, we perform a binary search for *i* in *Endpoints* to determine *t* and return *g* = *Strings*[*t, t* + *k*).

Lastly in this section, we point out that the minimizer length *m* controls a space/time trade-off for the proposed dictionary data structure. Small values of *m* create fewer and longer super-*k*-mers, thus lowering the space for the smaller values of *z* and *M*. On the other hand, *m* should not be chosen too small to avoid the scan of many super-*k*-mers at query time. We will experimentally show the trade-off in Section 5.1. Next, we take a deeper look at lookup time.

### 4.2 Skew Hashing

The efficiency of the Lookup query depends on the number of super-*k*-mers in the bucket of a minimizer, which we refer to as the “size” of the bucket. Since a minimizer can appear multiple times in the input strings, nothing prevents its bucket size to grow unbounded. For example, on the human genome, the largest bucket size can be as large as 3.6 × 10^4^ for *m* = 20 (or even larger for smaller values of *m*), meaning that a query inspecting such a bucket would be very slow in practice.

To avoid the burden of these heavy buckets, i.e., to guarantee that a Lookup inspects a *constant* number of buckets in the worst case, we exploit another important property of minimizers: the distribution of the bucket size is (very) *skewed* for sufficiently large *m*. That is, most minimizers appear just once and relatively few of them repeat many times – an observation also made in several previous works (see, e.g., [Chikhi et al., 2014, Jain et al., 2020].

Table 1 shows an example of such distribution for the first *n* = 10^9^ *k*-mers (for *k* = 31) of the human genome. Similar values were obtained for other genomes. (See the tables for other values of *n* in Supplementary Material). More precisely, a value in the table represents the fraction of buckets having size *s*, for *s* = 1, 2, 3, 4, 5 (only the first 5 sizes are shown for conciseness). The important thing to observe in the example is that, for *m* > 17, the distribution is very skewed, e.g., most buckets (> 90%) contain just 1 super-*k*-mer. As we want to take advantage of this distribution, we proceed as follows.

**Table 1.**
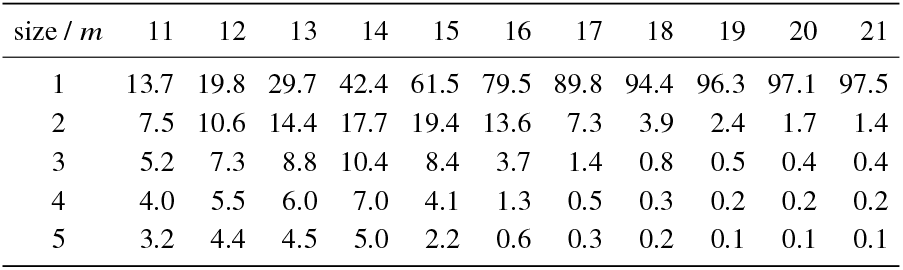
Bucket size distribution (%) for *k* = 31 and the first *n* = 10^9^ *k*-mers of the human genome, by varying minimizer length *m*.

We fix two quantities 𝓁 and *L*, with 0 ≤ 𝓁 < *L*. By virtue of the skew distribution, we have that the number of buckets whose size is larger than 2^𝓁^ is *small* for a proper choice of 𝓁, as well as the number of *k*-mers belonging to such buckets. For example with *m* = 20 and 𝓁 = 2, we see from Table 1 that we have 100.0 − (97.1 + 1.7 + 0.4 + 0.2) % = 0.6% of buckets with more than 2^𝓁=2^ = 4 super-*k*-mers. This allows us to build a minimal perfect hash function to speed up Lookup but *only for a small fraction* of the total *k*-mers – in marked contrast with prior schemes (reviewed in Section 3) that build the function over the entire set of *k*-mers. For ease of exposition, in the following we assume that 2𝓁 ≤ *max*, where *max* is the largest bucket size (the corner case for 2^𝓁^ ≤ *max* < 2^*L*^ is straightforward to handle). For 𝓁 ≤ *i* ≤ *L*, let 𝒮_*i*_ be the set of all the *k*-mers belonging to buckets of size *s*, with *s* such that

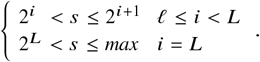

We build a MPHF *f*_*i*_ for each set 𝒮_*i*_. Now, given a *k*-mer *g* ∈ 𝒮_*i*_, we know that it belongs to a bucket containing at most 2^*i*+1^ super-*k*-mers. Therefore, we can store the identifier of the super-*k*-mer containing *g* in a vector, *V*_*i*_ [0, | 𝒮_*i*_ |), at position *f*_*i*_ (*g*). Importantly, each integer in *V*_*i*_ requires just *i* + 1 bits to be represented (*V*_*L*_ is formed by ⌈ log_2_ *max* ⌉ -bit integers). Fig. 2a illustrates this structure.

**Fig. 2.**
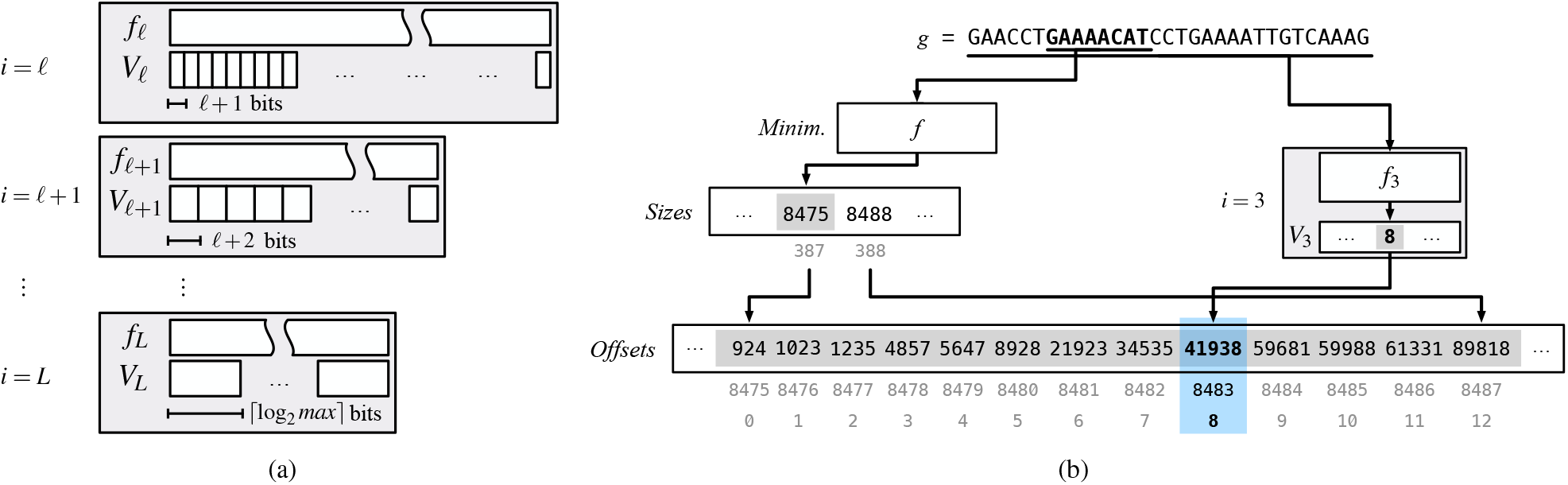
A schematic view of the skew index component of the dictionary (a), comprising partitions 𝓁 ≤ *i* ≤ *L*, each consisting of a MPHF *fi* and a compact vector *Vi*. Let us consider an example (b) of Lookup for *g* = GAACCT**GAAAACAT**CCTGAAAATTGTCAAAG and 𝓁 = 3. Suppose that the bucket for the minimizer *r* = GAAAACAT contains *s* = 13 super-*k*-mers (whose offsets are *Offsets*[8475, 8488) in the picture), thus it belongs to partition *i* = 3 because 2^3^ < 13 ≤ 2^3+1^. (Each integer in *V*_3_ is less than 2^3+1^, so it can be coded in *i* +1 = log_2_(2^3+1^) = 4 bits.) Now, also suppose that *g* is located in the 9-th super-*k*-mer of the bucket (i.e., that of index 8). It would then be time-consuming to fully scan the 8 super-*k*-mers before the 9-th. Therefore, we retrieve 8 – the index of the 9-th super-*k*-mer where *g* is located – from *V*3 [ *f*3 (*g*)] and know that *g* has to be searched for in the super-*k*-mer whose offset is *Offsets*[8475 + 8].

We point out that, again thanks to the skew distribution, it is very likely to have | 𝒮 | ≥ | 𝒮_+1_ | ≥ · · · ≥ | 𝒮_*L*−1_ |. Therefore, for a proper choice of 𝓁 and *L*, we expect this additional *skew index* component of the dictionary to take little space, while granting very fast searches.

To make a concrete example, let us consider the human genome and the skew index with 𝓁 = 6 and *L* = 12. So we form *L* − 𝓁 + 1 = 12 − 6 + 1 = 7 partitions; each partition is made up of a MPHF and a compact vector. Each MPHF *f*_*i*_ can be tuned to take 2.5 − 3.0 bits per key, whereas we spend *i* + 1 bits per integer in *V*_*i*_, *i* = 6, …, 11. (As already mentioned, *max* = 3.6 × 10^4^ for *m* = 20, thus we spend ⌈log_2_ *max* ⌉ = 16 bits per integer in *V*_*L*=12_.) The crucial point is that we have 0.016% of buckets that comprise more than 2^𝓁=6^ super-*k*-mers, for just 1.86% of the total *k*-mers. For this reason, the skew index costs less than 0.21 bits/*k*-mer over a total of 8.28 bits/*k*-mer (see also Fig. 1a of Supplementary Material).

#### Accelerated Lookup

Using the skew index to accelerate Lookup(*g*) is simple. As for regular Lookup, we compute the minimizer *r* of *g* and the quantities *begin* = *Sizes*[ *f* (*r*)] and *end* = *Sizes*[ *f* (*r*) + 1]. Therefore we know that the bucket of *r* has size *end* − *begin* ≤ *max*. Let *b* = ⌈log_2_(*end* − *begin*) ⌉. If *b* ≤ e, then the bucket is “small” and we proceed as already explained in Section 4.1. Otherwise we know that *g*, if present in the dictionary, belongs to some partition *i* of the skew index that, as per our description above, has MPHF *f*_*i*_ and compact vector *V*_*i*_ (*i* = *b* if *b* ≤ *L* or *i* = *L* otherwise). Thus, we retrieve the super-*k*-mer identifier *q* = *V*_*i*_ [ *f*_*i*_ (*g*)] and finally search for *g* in the super-*k*-mer whose offset is *Offsets*[*begin* + *q*]. (Note that if *q* ≥ *end*, then *g* cannot belong to the dictionary.) In conclusion, although the skew index performs 2 additional accesses per Lookup, one for *f*_*i*_ and one for *V*_*i*_, it limits the number of accesses made to *Strings* to 2. (To handle reverse complements, we may have to repeat the process also for the reverse complement of *g*.)

Fig. 2b shows a concrete example of Lookup.

### 4.3 Streaming Queries

The Lookup algorithm we have described in the previous section is *contextless*, i.e., it does not take advantage of the specific, *consecutive*, query order issued by sequence analysis tasks. As already mentioned in Section 1, given a string *P* of length | *P* | ≥ *k*, we are interested in determining the result of Lookup for all the *k*-mers read consecutively from *P*. We would like to do it faster than just performing | *P* | − *k* + 1 independent lookups. Therefore, in this section we describe some important optimizations for streaming lookup queries that work well with the proposed dictionary data structure. The general idea is to cache some extra information about the result for the *k*-mer *g* = *P*[*i, i* + *k*) to speed up the computation for the next *k*-mer in *P*, say *g*_*nx*_ = *P*[*i* + 1, *i* + *k* + 1).

The algorithm keeps track of the minimizer *r* of *g* and the position *j* at which the last match was found in *Strings*, i.e., if *g* belongs to the dictionary, then it is located at *Strings*[*j, j* + *k*) for some *j*. These two variables make up a *state* information that is updated during the execution of the algorithm. Given that consecutive *k*-mers are likely to share the same minimizers, we compare *r* to the minimizer of *g*_*nx*_, say *r*_*nx*_.

- If *r*_*nx*_ = *r*, then we know that *g*_*nx*_ belongs to the same bucket *B*_*r*_ of *g*, thus we avoid recomputing *f* and spare the accesses to both *Sizes* and *Offsets*. Also, if *g* was actually found in the dictionary (therefore, starting at *Strings*[*j*]) good chances are that *g*_*nx*_ is found at *Strings*[*j* + 1]. If so, we refer to the latter matching case as an *extension*. Intuitively, if the algorithm “extends” frequently, i.e., most matches in *P* are determined by just looking at consecutive *k*-mers in *Strings*, then fast evaluation is retained. If the algorithm does not extend from *g* to *g*_*nx*_, i.e., *g*_*nx*_ is not found at *Strings*[*j* + 1], then we scan the bucket *B*_*r*_. Therefore, if present in the dictionary, *g*_*nx*_ will be found at some other position *j*_*nx*_. So we update the state by setting *j* = *j*_*nx*_.
- If *r*_*nx*_ * *r*, then we proceed as for a regular Lookup query, locating the new bucket *B*_*r*_*nx* and searching for *g*_*nx*_. We then set *r* = *r*_*nx*_.

Of course it can happen that the minimizer *r* does not belong to the set of minimizers indexed by the dictionary. Recall from Section 4.1 that we build the MPHF *f* for the set M of all the distinct minimizers in the input. In this case, we are sure that any *k*-mer *g* whose minimizer *r* < ℳ is not to be found in the dictionary. By definition, however, we are not able to detect if *r* < M using the MPHF *f*. That is, *f* will still locate a bucket and all the *k*-mers in the bucket will have (the same) minimizer, different from *r*. Therefore, when searching for *g*, we first compare *r* with the minimizer of the first *k*-mer read in the bucket: if they are different, we know that *r* < ℳ and *g* does not belong to the dictionary. In the case when *r* < ℳ, the algorithm still caches the last seen minimizer because if *r*_*nx*_ = *r* then also *r*_*nx*_ < ℳ and *g*_*nx*_ cannot belong to the dictionary.

In conclusion – as long as the minimizer is the same – either the algorithm works *locally* in the same bucket, or safely skips the search.

Another convenient information to cache in the state of the algorithm is the *orientation* of the last match, that is, whether the last queried *k*-mer *g* was found in the dictionary as *g* or as its reverse complement *ĝ*. In fact, if *g* was found as *g* then also *g*_*nx*_ is likely to be found as *g*_*nx*_ and extension should be tried in *forward* direction (say, from lower to higher offsets in *Strings*). But if *g* was found as *ĝ*, then is more efficient to try to extend the matching in *backward* direction, hence effectively iterating backwards in *Strings*. In fact, suppose that the whole string *P* (for ease of exposition) is present in *Strings* but in its reverse complement form. Then the first *k*-mer *g* of *P* will be found as *ĝ* in last position in the located “region” of *Strings*, say at some position *j*. Any other attempt to extend the matching in forward direction (from *j* to *j* + 1) will then fail and any subsequent *g*_*nx*_ will be searched for by re-scanning the bucket again. That is, we end up in scanning the bucket for *c* = | *P* | − *k* + 1 times, for at least *O*(*c*^2^) *k*-mer comparisons. To prevent this quadratic behavior in case of reverse complemented patterns, we try to directly extend the matching for *g*_*nx*_ moving from *j* to *j* − 1.

## 5 Experiments

In this section we benchmark the proposed dictionary data structure – which we refer to as SSHash in the following – and compare it against the indexes reviewed in Section 3. For all our experiments, we fix *k* to 31.

Our implementation of SSHash is written in C++17 and available at https://github.com/jermp/sshash. For the experiments we report here, the code was compiled with gcc 11.2.0 under Ubuntu 19.10 (Linux kernel 5.3.0, 64 bits), using the flags -O3 and -march=native. We do not explicitly use any SIMD instruction in our codebase.

We use a server machine equipped with an Intel i9-9940X processor (clocked at 3.30 GHz) and 128 GB of RAM. The reported timings were collected using a single core of the processor. All dictionaries were fully loaded in internal memory before running the experiments. The SSHash dictionaries were also built entirely in internal memory.

### Datasets

We downloaded some DNA collections (in .fasta format) and built the compacted de Bruijn graph using the tool BCALM (v2) [Chikhi et al., 2016], without any *k*-mer filtering, to extract the maximal unitigs. We then run the tool UST [Rahman and Medvedev, 2020] to compute the corresponding path covers. Table 2 reports the basic statistics of the path covers. In particular we used: the whole genomes of the atlantic cod (*Gadus Morhua*) and the common kestrel (*Falco Tinnunculus*), the whole GRCh38 human genome (*Homo Sapiens*), and a collection of more than 8000 bacterial genomes from Almodaresi et al. [2018].

**Table 2.**
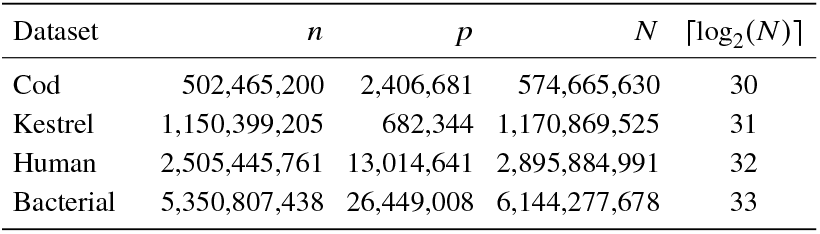
Some basic statistics for the datasets used in the experiments, for *k* = 31, such as number of: *k*-mers (*n*), paths (*p*), and bases (*N*).

| Dataset | $n$ | $p$ | $N$ | $\lceil \log_2(N) \rceil$ |
| --- | --- | --- | --- | --- |
| Cod | 502,465,200 | 2,406,681 | 574,665,630 | 30 |
| Kestrel | 1,150,399,205 | 682,344 | 1,170,869,525 | 31 |
| Human | 2,505,445,761 | 13,014,641 | 2,895,884,991 | 32 |
| Bacterial | 5,350,807,438 | 26,449,008 | 6,144,277,678 | 33 |

At the code repository https://github.com/jermp/sshash we provide further instructions on how to download and prepare the datasets for indexing.

#### 5.1 Tuning

Before comparing SSHash against other dictionaries, we first benchmark SSHash in isolation to fix a suitable choice for *m* and quantify the impact of the different parsing modalities (regular vs. canonical) that we introduced in Section 4.1. Following our discussion in Section 4.2, we use 𝓁 = 6 and *L* = 12 for all SSHash dictionaries.

To measure query time, we use 10^6^ queries and report the mean between 5 measurements. For positive lookups, i.e., those for *k*-mers present in the dictionary, we sampled uniformly at random 10^6^ *k*-mers from each collection and use them as queries. Very importantly, 50% of them were transformed into their reverse complements to make sure we benchmark the dictionaries in the most general case. For negative lookups, we simply use randomly generated *k*-mer strings. For Access, we generated 10^6^ integers uniformly at random in the range [0, *n*) for each collection and extract the corresponding *k*-mer strings.

#### Access and Iteration Time

We first recall that the time for Access (and thus, that for iteration) does not depend on *m* nor 𝓁. The average Access time is, instead, affected by the size of the data structure, i.e., by *n* and *p*: Access is on average 2 − 3× faster than Lookup since the wanted string is accessed directly, rather than searched for in the dictionary. Iterating thorough all *k*-mers in the dictionary is very fast and even independent from *n*: on average, it costs 20 − 22 ns/*k*-mer. Therefore, for the rest of this section we entirely focus on lookup time.

#### Space and Lookup Time

With the help of Table 1 in Supplementary Material (see also the example at page 5 for *n* = 10^9^), we choose some suitable ranges of *m* for the different dataset sizes. The space/time trade-off by varying *m* in such ranges, for both regular and canonical parsing modalities, is shown in Table 3. As we discussed in Section 4.1 and apparent from the table, *m* controls a trade-off between dictionary size and lookup time: the smaller the *m* value, the more compact the dictionary, but the slower the dictionary as well (and vice versa). While it is difficult to precisely tell by how much the space will grow when moving from *m* to *m* + 1, we see that the space grows by ≈ 0.3 − 0.4 bits/*k*-mer, for both regular and canonical parsing. The canonical parsing modality costs ≈ 1.0 − 1.5 bits/*k*-mer more than the regular one for the same value of *m* because more distinct minimizers are used. However, the canonical version improves lookup time significantly (especially for negative queries), by a factor of 1.4 − 2.0× on average, because only one bucket per query is inspected in the worst case rather than two by the regular modality.

**Table 3.** Space in bits/*k*-mer (bpk) and Lookup time (indicated by Lkp^+^ for positive queries; by Lkp^−^ for negative) in average ns/*k*-mer for regular and canonical SSHash dictionaries by varying minimizer length *m*. For each dataset, we indicate promising configurations in bold font.

| Dataset | $m$ | | | $m$ | | | $m$ | | | $m$ | | |
| --- | --- | --- | --- | --- | --- | --- | --- | --- | --- | --- | --- | --- |
| | bpk | $Lkp^+$ | $Lkp^-$ | bpk | $Lkp^+$ | $Lkp^-$ | bpk | $Lkp^+$ | $Lkp^-$ | bpk | $Lkp^+$ | $Lkp^-$ |
| Cod | 15 |  |  | **16** |  |  | **17** |  |  | 18 |  |  |
| regular | 6.60 | 1236 | 1267 | 6.82 | 1100 | 1174 | **6.98** | **1045** | **1158** | 7.21 | 1015 | 1157 |
| canonical | 7.68 | 945 | 768 | **7.92** | **834** | **690** | 8.18 | 786 | 672 | 8.47 | 755 | 658 |
| Kestrel | **16** |  |  | **17** |  |  | 18 |  |  | 19 |  |  |
| regular | 6.19 | 1137 | 1323 | **6.48** | **1042** | **1265** | 6.79 | 1005 | 1245 | 7.12 | 997 | 1240 |
| canonical | **7.30** | **882** | **781** | 7.68 | 790 | 722 | 8.09 | 743 | 696 | 8.51 | 730 | 691 |
| Human | 17 |  |  | 18 |  |  | **19** |  |  | **20** |  |  |
| regular | 7.44 | 1591 | 1668 | 7.67 | 1459 | 1573 | 7.95 | 1406 | 1547 | **8.28** | **1338** | **1530** |
| canonical | 8.76 | 1150 | 936 | 9.04 | 1054 | 881 | **9.39** | **990** | **854** | 9.80 | 958 | 838 |
| Bacterial | 18 |  |  | **19** |  |  | **20** |  |  | 21 |  |  |
| regular | 7.42 | 1535 | 1867 | 7.80 | 1425 | 1813 | **8.22** | **1389** | **1780** | 8.70 | 1368 | 1774 |
| canonical | 8.75 | 1129 | 1043 | **9.22** | **1051** | **995** | 9.75 | 1028 | 947 | 10.34 | 998 | 956 |

Since we seek for a good balance between dictionary space and lookup time, in the light of the results reported in Table 3, we choose *m* as follows. For Cod and Kestrel: *m* = 17 with regular parsing; *m* = 16 with canonical parsing; for Human and Bacterial: *m* = 20 with regular parsing; *m* = 19 with canonical parsing. In the following, we assume these values of *m* are used and omit the indication from the tables.

In general, we observe that a good value for *m* satisfies 4^*m*^ > *N*, i.e., *m* should be chosen as to have – at least – as many possible minimizers as the number of bases in the input. It is therefore recommended to use *m* = ⌈log_4_(*N*) ⌉ + 1 or *m* = ⌈log_4_(*N*) ⌉ + 2.

### 5.2 Comparison Against Other Dictionaries

In this section we compare SSHash against the following state-of-the-art dictionaries that we briefly reviewed in Section 3:

- dBG-FM [Chikhi et al., 2014] – An implementation of the popular FM-index tailored for DNA. This implementation is widely used as an exact membership data structure for *k*-mers [Chikhi et al., 2014, Rahman and Medvedev, 2020], also in the ABySS assembler [Simpson et al., 2009, Jackman et al., 2017]. We tested the index by sampling one position of the suffix-array every *s* positions, for *s* = 32, 64, 128.
- Pufferfish [Almodaresi et al., 2018] – We test both the dense and sparse versions of the index. The sparse version was obtained with parameters *s* = 9 and *e* = 4 as used in the original paper.
- Blight [Marchet et al., 2021] – We test the index with sampling rate *b* = 0, 2, 4 and minimizer length *m* = 10 as suggested in the paper. We recall that a sampling rate of *b* > 0 reduces the index space by *b* bits/*k*-mer at the expense of query time.

We use the C++ implementations from the respective authors: links to the various GitHub libraries are provided in the References. All sources were compiled using the same compilation flags as used for SSHash.

#### Space and Lookup Time

We first consider the space of the dictionaries, reported in Table 4. The space of SSHash is significantly better than that of the other approaches based on minimal perfect hashing, roughly: 2 − 2.5× (or more) better than Blight, and 3 − 5× better than Pufferfish. This is so primarily because these approaches build a MPHF for the *entire* set of *k*-mers, hence associate a positional information (e.g., in the reference genome) to *each k*-mer in the input. We point out that, unlike for Blight, this is expected for Pufferfish dense since it was exactly designed for the purpose of reference mapping. (The shaded rows in Table 4 account for the space needed by Pufferfish to only support Lookup, i.e., discarding the color information in its colored de Bruijn graph structure.)

**Table 4.** Dictionary space in total GB and average bits/*k*-mer (bpk).

| Dictionary | Cod |  | Kestrel |  | Human |  | Bacterial |  |
| --- | --- | --- | --- | --- | --- | --- | --- | --- |
|  | GB | bpk | GB | bpk | GB | bpk | GB | bpk |
| dBG-FM, $s = 128$ | 0.22 | 3.48 | 0.44 | 3.07 | – | – | – | – |
| dBG-FM, $s = 64$ | 0.27 | 4.38 | 0.55 | 3.86 | – | – | – | – |
| dBG-FM, $s = 32$ | 0.39 | 6.16 | 0.78 | 5.43 | – | – | – | – |
| Pufferfish, sparse | 1.75 | 27.80 | 3.69 | 25.66 | 8.87 | 28.32 | 18.91 | 28.28 |
|  | 1.49 | 23.70 | 3.37 | 23.40 | 7.50 | 23.96 | 16.09 | 24.06 |
| Pufferfish, dense | 2.69 | 42.76 | 5.97 | 41.54 | 14.11 | 45.04 | 30.70 | 45.89 |
|  | 2.43 | 38.66 | 5.65 | 39.28 | 12.74 | 40.68 | 27.88 | 41.68 |
| Blight, $b = 4$ | 0.91 | 14.53 | 2.16 | 15.00 | 5.04 | 16.11 | 11.40 | 17.04 |
| Blight, $b = 2$ | 1.04 | 16.57 | 2.45 | 17.04 | 5.67 | 18.13 | 12.74 | 19.05 |
| Blight, $b = 0$ | 1.17 | 18.61 | 2.74 | 19.06 | 6.32 | 20.17 | 14.12 | 21.11 |
| SSHash, regular | 0.44 | 6.98 | 0.93 | 6.48 | 2.59 | 8.28 | 5.50 | 8.22 |
| SSHash, canonical | 0.50 | 7.92 | 1.00 | 7.30 | 2.94 | 9.39 | 6.17 | 9.22 |

The dBG-FM index is, not surprisingly, the most compact, thanks to the compression of the powerful Burrows-Wheeler transform (BWT) [Burrows and Wheeler, 1994]. (We were unable to build the index correctly on the larger Human and Bacterial datasets.) While dBG-FM is several times smaller than Pufferfish and Blight, note that its smallest version tested (for *s* = 128) is only essentially 2× smaller than regular SSHash and this gap diminishes at higher sampling rates. For example, dBG-FM for *s* = 32 is only 13 − 17% smaller than regular SSHash. However, SSHash answers lookup queries *much* faster than dBG-FM as shown in Table 5. A lookup query in the dBG-FM index is implemented as a classic *count* query on a FM-index (see the paper by Ferragina and Manzini [2000] for details) which, for a pattern of length *k*, generates at least *k* cache-misses. This cost is even higher for the handling of reverse complements that may induce two distinct count queries.

**Table 5.**
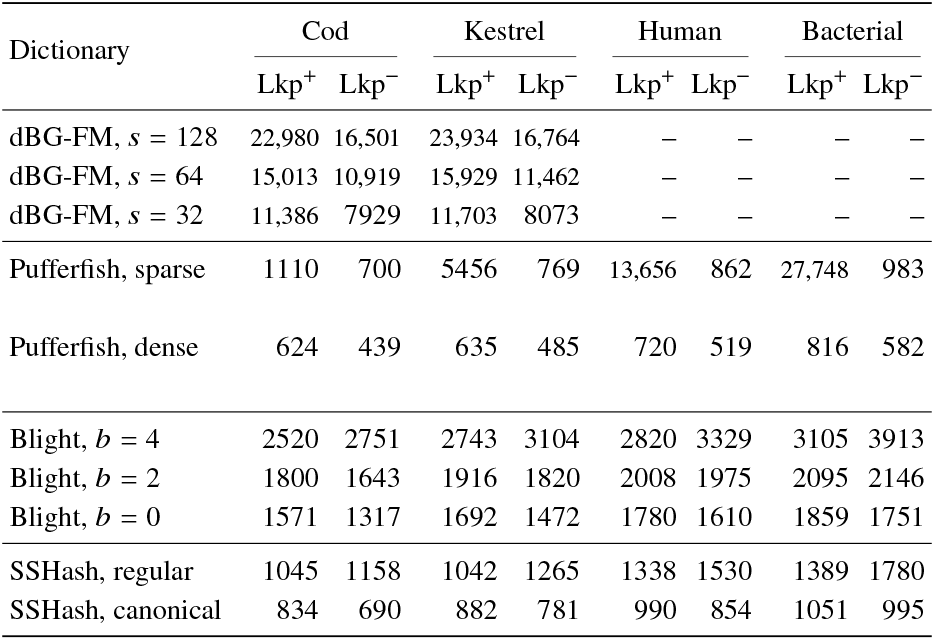
Dictionary Lookup time in average ns/*k*-mer.

| Dictionary | Cod |  | Kestrel |  | Human |  | Bacterial |  |
| --- | --- | --- | --- | --- | --- | --- | --- | --- |
|  | Lkp<sup>+</sup> | Lkp<sup>−</sup> | Lkp<sup>+</sup> | Lkp<sup>−</sup> | Lkp<sup>+</sup> | Lkp<sup>−</sup> | Lkp<sup>+</sup> | Lkp<sup>−</sup> |
| dBG-FM, $s = 128$ | 22,980 | 16,501 | 23,934 | 16,764 | – | – | – | – |
| dBG-FM, $s = 64$ | 15,013 | 10,919 | 15,929 | 11,462 | – | – | – | – |
| dBG-FM, $s = 32$ | 11,386 | 7929 | 11,703 | 8073 | – | – | – | – |
| Pufferfish, sparse | 1110 | 700 | 5456 | 769 | 13,656 | 862 | 27,748 | 983 |
| Pufferfish, dense | 624 | 439 | 635 | 485 | 720 | 519 | 816 | 582 |
| Blight, $b = 4$ | 2520 | 2751 | 2743 | 3104 | 2820 | 3329 | 3105 | 3913 |
| Blight, $b = 2$ | 1800 | 1643 | 1916 | 1820 | 2008 | 1975 | 2095 | 2146 |
| Blight, $b = 0$ | 1571 | 1317 | 1692 | 1472 | 1780 | 1610 | 1859 | 1751 |
| SSHash, regular | 1045 | 1158 | 1042 | 1265 | 1338 | 1530 | 1389 | 1780 |
| SSHash, canonical | 834 | 690 | 882 | 781 | 990 | 854 | 1051 | 995 |

We observe that SSHash regular is as fast as (or faster than) the fastest Blight’s version, for *b* = 0, and faster for higher *b*. SSHash canonical is always much faster than Blight. Pufferfish dense is instead faster than SSHash thanks to its simpler lookup procedure that just needs to retrieve the absolute offset of a *k*-mer using hashing and check the *k*-mer against the reference string. However, we point out that: (1) this higher Lookup efficiency comes at a significant penalty in space effectiveness compared to SSHash, and (2) the sparse variant’s performance degrades on larger datasets.

#### Streaming Membership Query Time

We now consider streaming membership queries. Pufferfish and Blight are also optimized to answer these kind of stateful queries. To query the dictionaries, we use some reads (in .fastq format) downloaded from the European Nucleotide Archive (ENA), and related to each dataset – Cod: run accession SRR12858649 with 2,041,092 reads, each of length 110 bases; Kestrel: run accession SRR11449743 with 14,647,106 reads, each of length 125 bases; Human: run accession SRR5833294 with 34,129,891 reads, each of length 76 bases; Bacterial: run accession SRR5901135 with 4,628,576 reads of variable length (a sequencing run of *Escherichia Coli*).

We lookup for every *k*-mer read in sequence from the query files. For all the indexes, we just count the number of returned results rather than saving them to a vector. The result is reported in Table 6.

**Table 6.**
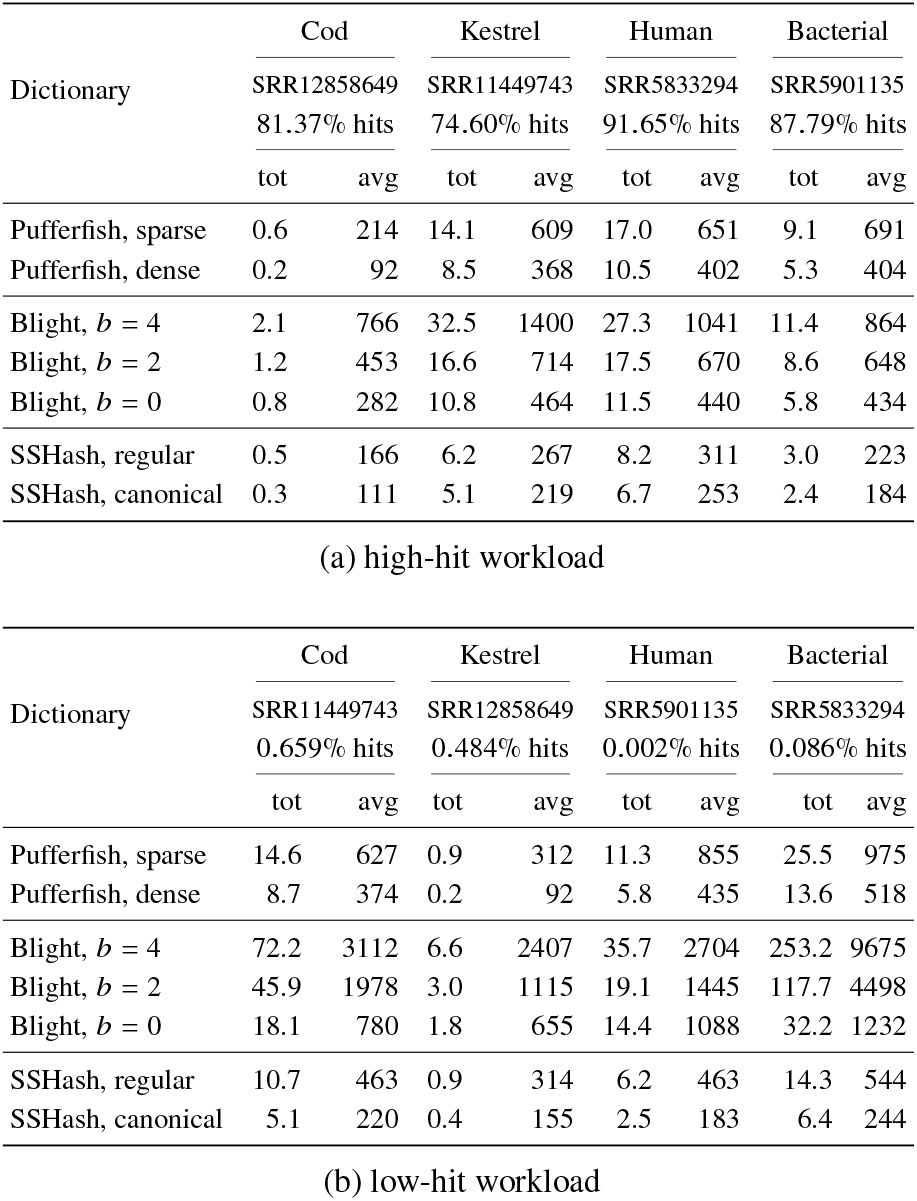
Query time for streaming membership queries for various dictionaries. The query time is reported as total time in minutes (tot), and average ns/*k*-mer (avg). We also indicate the query file (SRR number) and the percentage of hits. Both high-hit (> 70% hits) and low-hit (< 1% hits) workloads are considered.

In general terms, we see that SSHash is either comparable to or faster (by 2 − 3×) than Pufferfish and Blight. This holds true for both high-hit workloads (> 70% hits, i.e., *k*-mers present in the dictionary) and low-hit workloads (< 1% hits). It is important to benchmark the dictionaries under these two different query scenarios as both situations are meaningful in practice. (In our experiments, low-hit workloads are obtained by querying the dictionaries using a different query file as indicated in Table 6.) Indeed, observe that while Pufferfish’s performance is robust under both scenarios, Blight’s query time significantly degrades when most queries are negative, especially for *b* > 0. Also regular SSHash is almost 2× slower for low-hit workloads compared to high-hit workloads. This is expected, however, because almost all queries are exhaustively inspecting two buckets per *k*-mer as we explained in Section 4.1. Note that its performance is anyway better than Blight’s and not much worse than Pufferfish’s (dense variant). The canonical version of SSHash protects against this behavior in case of low-hit workload and, in fact, is generally the fastest dictionary.

Another meaningful point to mention is that SSHash does not allocate extra memory at query time, i.e., only the memory of the index – as reported in Table 4 – is retained (the memory for the state information maintained by the streaming algorithm described in Section 4.3 is constant). Pufferfish also does not allocate extra memory. Blight, instead, consumes more memory at query time than that required by its index layout on disk. For example, to perform the queries on the Human dataset in Table 6a, Blight with *b* = 0 uses a maximum resident set size of 7.51 GB compared to the 6.32 GB taken by its index on disk (23.98 vs. 20.17 bits/*k*-mer). This effect is even accentuated for higher *b* values.

#### Construction Time

Table 7 reports the time and internal memory used to build the dictionaries. The dBG-FM index needs to build the BWT of the input prior to indexing. This step can be very time consuming for large collections such as the ones of practical interest. That is, another important advantage of schemes based on hashing compared to BWT-based indexes is that they require significantly less time to build. This is evident from the result reported in the table.

**Table 7.** Dictionary construction times in minutes (using a single processing thread) and peak internal memory used during construction in GB. (Blight’s performance was the same for all values of *b* in the experiment.)

| Dictionary | Cod |  | Kestrel |  | Human |  | Bacterial |  |
| --- | --- | --- | --- | --- | --- | --- | --- | --- |
|  | min | GB | min | GB | min | GB | min | GB |
| dBGM-FM, $s = 128$ | 28.5 | 0.5 | 100.0 | 0.7 | – | – | – | – |
| dBGM-FM, $s = 64$ | 28.5 | 0.6 | 100.0 | 0.9 | – | – | – | – |
| dBGM-FM, $s = 32$ | 28.5 | 0.7 | 100.0 | 1.1 | – | – | – | – |
| Pufferfish, sparse | 15.5 | 3.3 | 35.2 | 6.7 | 86.0 | 19.4 | 200.8 | 40.1 |
| Pufferfish, dense | 13.0 | 2.8 | 29.2 | 5.9 | 70.7 | 14.0 | 173.2 | 30.4 |
| Blight | 5.0 | 3.3 | 11.0 | 7.0 | 25.0 | 7.5 | 50.0 | 15.8 |
| SSHash, regular | 1.5 | 2.6 | 3.8 | 5.7 | 12.5 | 15.4 | 29.6 | 33.4 |
| SSHash, canonical | 2.0 | 2.8 | 4.4 | 5.8 | 16.2 | 17.3 | 36.0 | 36.6 |

Due to space constraints, we do not describe the SSHash’s construction algorithm in this article. We just point out that the construction is efficient; indeed SSHash took much less time to build on the test collections compared to both Blight and Pufferfish. Its memory usage is comparable to that of Pufferfish, while Blight scales better in this regard (e.g., by retaining 2× less internal memory on Human and Bacterial) as it partially uses external memory. In future work, we will adapt the SSHash dictionary construction to use external memory too.

We also note that the SSHash canonical takes consistently more time and space to build than the regular variant: this is a direct consequence of the denser sampling of minimizers.

## 6 Conclusions and Future Work

We have studied the compressed dictionary problem for *k*-mers and proposed a solution, SSHash, based on a careful orchestration of minimal perfect hashing and compact encodings. In particular, SSHash is an exact and associative *k*-mer dictionary designed to deliver good practical performance. From a technical perspective, SSHash exploits the sparseness and the skew distribution of *k*-mer minimizers to achieve compact space, while allowing fast lookup queries.

We tested SSHash on collections of billions of *k*-mers and compared it against other indexes, under different query workloads (high-vs. low-hit) and modalities (random vs. streaming). Our implementation of SSHash is written in C++ and open source.

Compared to BWT-based indexes (like the dBG-FM index), SSHash is more than one order of magnitudes faster at lookup for only 2× larger space on average. Compared to prior schemes based on minimal perfect hashing (like Pufferfish and Blight), SSHash is significantly more compact (2 − 5× depending on the configuration) without sacrificing query efficiency. Indeed, SSHash is also the fastest dictionary for streaming membership queries. For these reasons we believe that SSHash embodies a superior space/time trade-off for the problem tackled in this work.

Several avenues for future work are possible. We mention some promising ones. First, we will engineer the dictionary construction to use multi-threading and external memory. Parallel query processing is also interesting; since SSHash is a read-only data structure, its queries are amenable to parallelism. We could also add support for other types of queries, such as *navigational* queries [Chikhi et al., 2014] that, given a *k*-mer *g*, ask to enumerate *all* the extensions of *g* (i.e., in both forward and backward direction) that are present in the dictionary. Another promising direction could adapt the SSHash data structure to also store the abundances of *k*-mers, which is a separate but related problem in the literature [Shibuya et al., 2021, Italiano et al., 2021]. Based on the observation that consecutive *k*-mers tend to have the same or very similar abundance we expect to add a small extra space to SSHash to store this information. In this paper we focused on minimizers for their simplicity and practical efficiency but one could also explore the effects of replacing the minimizers with other types of string sampling mechanisms [Loukides and Pissis, 2021, Sahlin, 2021]. Lastly, we also plan to study the *approximate* version of the dictioanary problem where it is allowed to tolerate a prescribed false positive rate.

## Acknowledgments

We thank Rossano Venturini for useful discussions on this work. We also thank the reviewers for their valuable suggestions that led to an improved presentation of the article.

