## Supplementary Material for "Sparse and Skew Hashing of K-Mers"

OXFORD

### Abstract

This document contains the Supplementary Material for the paper "Sparse and Skew Hashing of K-Mers".
**Contact:**

### Window Size during Lookup Queries

Section 4.1 contains a description of the Lookup algorithm. Here, we provide a further detail.

By design, we cannot compute the length of a super-$k$-mer from the *Offsets* array (at least, not efficiently). That is, the offset $t$ only indicates that a super-$k$-mer *begins* at *Strings*$[t]$. So we start comparing the $k$-mer at *Strings*$[t]$ but we do not know exactly after how many symbols we can stop the search. We could re-compute minimizers from the $k$-mers read during the scan of *Strings* to derive this information but we want to avoid doing so. In fact, we know that a super-$k$-mer $g'$ cannot contain more than $k - m + 1$ $k$-mers unless the *same* minimizer appears again somewhere in $g'[0, 2k - m)$. Albeit rare, the latter case is possible and it is handled as follows. If a super-$k$-mer is longer than $2k - m$ symbols, it is split into blocks of $2k - m$ symbols each (except possibly the last one) and we just pretend that every block is a distinct super-$k$-mer. Thus, at query time it is safe to consider a window of size $W = \min(2k - m, t_{end} - t)$ symbols, where $t_{end} \geq t + k$ is the *successor* of the offset $t$ in *Endpoints* (i.e., $t_{end}$ is the smallest value in *Endpoints* that is lager-than or equal-to $t$). Capping $W$ to $t_{end} - t$ symbols if $t_{end} - t < 2k - m$ is necessary to not falsely consider alien $k$-mers that are formed at the boundary between two consecutive strings in the compact vector *Strings*.

### Bucket Size Distribution

Table 1 shows an example of the skew distribution of bucket size, for some indicative values of $n$ (these values mimic the ones used in the experiments in Section 5). More precisely, a value in a table represents the fraction of buckets having size $s$ (number of super-$k$-mers), for $s = 1, 2, 3, 4, 5$ (only the first 5 sizes are shown for conciseness).

### Space Breakdowns

It is also interesting to report how the space is subdivided into the different dictionary components. Fig. 1 shows an example of space breakdown for both regular and canonical dictionaries built from the Human dataset. Very similar breakdowns are obtained for the other datasets. As expected, the most space-consuming component in the dictionary is the *Offsets* component (61 − 63% of the total space). Hence, any effort to make SSHash more compact should be spent in making this component more light-weight. In this regard we recall that, because of the skew distribution of bucket size, most of the offsets' space is due to buckets of size 1: hence, *Offsets* is an array made by integers in the range $[0, N)$ essentially shuffled at random by the minimizers' MPHF. Therefore, our choice of spending $\lceil \log_2(N) \rceil$ bits per offset is basically optimal for such distribution. Lastly, the space difference between a regular and a canonical dictionary almost entirely resides in the *Offsets* component given that more minimizers are used, thus realizing a denser sampling.

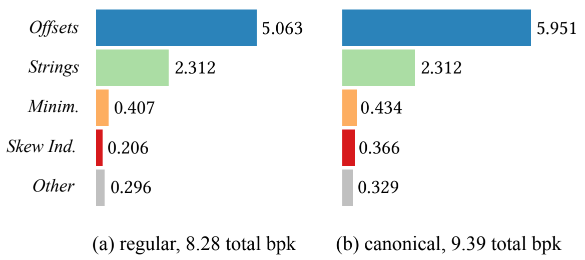


**Fig. 1.** Space breakdowns for the Human dataset, for both (a) regular and (b) canonical dictionaries. The numbers next to each bar indicate the bits/$k$-mer (bpk) spent by the respective components.

The second most space-consuming component is *Strings* (25 − 28% of the total space), which costs $2N/n$ bits/$k$-mer regardless of the parsing modality. While it could be possible to obtain better compression for *Strings* using a compressor for DNA (we only need to sequentially decompress a super-$k$-mer at lookup time) [Manzini and Rastero, 2004], we do not consider this as a promising option to explore given that most of the dictionary space is spent elsewhere (and given the slowdown in query processing that would follow).

Both the MPHF on the minimizers (*Minim.*) and the skew index take a small fraction of the total space (e.g., 5% and 2.5% respectively for the regular variant, Fig. 1a). Other costs include those for representing the endpoints of the paths (*Endpoints* array) and size of the buckets (*Sizes* array): these costs are small, especially thanks to the use of the Elias-Fano encoding.

© The Author 2022.

Table 1. Bucket size distribution (%) for $k = 31$ and some useful values of $n$ (number of $k$-mers), by varying $m$ (minimizer length).

| size / $m$ | 11 | 12 | 13 | 14 | 15 | 16 | 17 | 18 | 19 | 20 | 21 |
|---|---|---|---|---|---|---|---|---|---|---|---|
| 1 | 16.1 | 24.0 | 35.3 | 51.3 | 71.6 | 85.9 | 92.8 | 95.6 | 96.8 | 97.4 | 97.7 |
| 2 | 8.7 | 12.6 | 15.7 | 19.7 | 17.0 | 10.0 | 5.2 | 2.9 | 2.0 | 1.6 | 1.4 |
| 3 | 6.1 | 8.3 | 9.4 | 10.4 | 5.7 | 2.2 | 1.0 | 0.6 | 0.5 | 0.4 | 0.4 |
| 4 | 4.6 | 6.0 | 6.6 | 6.1 | 2.3 | 0.8 | 0.4 | 0.3 | 0.2 | 0.2 | 0.2 |
| 5 | 3.8 | 4.6 | 5.1 | 3.7 | 1.1 | 0.4 | 0.2 | 0.1 | 0.1 | 0.1 | 0.1 |

(a) $n = 0.5 \times 10^9$

| size / $m$ | 11 | 12 | 13 | 14 | 15 | 16 | 17 | 18 | 19 | 20 | 21 |
|---|---|---|---|---|---|---|---|---|---|---|---|
| 1 | 13.7 | 19.8 | 29.7 | 42.4 | 61.5 | 79.5 | 89.8 | 94.4 | 96.3 | 97.1 | 97.5 |
| 2 | 7.5 | 10.6 | 14.4 | 17.7 | 19.4 | 13.6 | 7.3 | 3.9 | 2.4 | 1.7 | 1.4 |
| 3 | 5.2 | 7.3 | 8.8 | 10.4 | 8.4 | 3.7 | 1.4 | 0.8 | 0.5 | 0.4 | 0.4 |
| 4 | 4.0 | 5.5 | 6.0 | 7.0 | 4.1 | 1.3 | 0.5 | 0.3 | 0.2 | 0.2 | 0.2 |
| 5 | 3.2 | 4.4 | 4.5 | 5.0 | 2.2 | 0.6 | 0.3 | 0.2 | 0.1 | 0.1 | 0.1 |

(b) $n = 1.0 \times 10^9$

| size / $m$ | 11 | 12 | 13 | 14 | 15 | 16 | 17 | 18 | 19 | 20 | 21 |
|---|---|---|---|---|---|---|---|---|---|---|---|
| 1 | 11.2 | 15.6 | 23.0 | 33.9 | 48.2 | 67.8 | 83.2 | 91.3 | 94.9 | 96.4 | 97.1 |
| 2 | 6.1 | 8.5 | 12.1 | 15.3 | 18.9 | 18.0 | 11.4 | 6.2 | 3.5 | 2.3 | 1.8 |
| 3 | 4.3 | 5.9 | 8.1 | 9.1 | 10.4 | 6.7 | 2.8 | 1.2 | 0.7 | 0.5 | 0.4 |
| 4 | 3.3 | 4.5 | 5.9 | 6.3 | 6.5 | 3.0 | 1.0 | 0.4 | 0.3 | 0.2 | 0.2 |
| 5 | 2.7 | 3.7 | 4.6 | 4.8 | 4.2 | 1.5 | 0.5 | 0.2 | 0.2 | 0.1 | 0.1 |

(c) $n = 2.5 \times 10^9$

| size / $m$ | 11 | 12 | 13 | 14 | 15 | 16 | 17 | 18 | 19 | 20 | 21 |
|---|---|---|---|---|---|---|---|---|---|---|---|
| 1 | 8.7 | 11.2 | 15.1 | 23.6 | 41.4 | 62.9 | 78.2 | 86.9 | 91.6 | 94.1 | 95.4 |
| 2 | 4.8 | 6.1 | 8.4 | 13.5 | 19.9 | 19.1 | 13.7 | 9.2 | 6.4 | 4.7 | 3.8 |
| 3 | 3.3 | 4.2 | 6.0 | 9.6 | 11.5 | 7.7 | 4.1 | 2.2 | 1.3 | 0.8 | 0.6 |
| 4 | 2.5 | 3.3 | 4.7 | 7.3 | 7.2 | 3.7 | 1.6 | 0.8 | 0.4 | 0.2 | 0.1 |
| 5 | 2.1 | 2.7 | 3.9 | 5.9 | 4.8 | 2.0 | 0.8 | 0.4 | 0.2 | 0.1 | 0.1 |

(d) $n = 5.0 \times 10^9$
